# Four novel, immortal cell lines derived from embryos of *Xenopus tropicalis*

**DOI:** 10.1101/2022.03.01.482505

**Authors:** Gary J. Gorbsky, John R. Daum, Hem Sapkota, Katja Summala, Hitoshi Yoshida, Constantin Georgescu, Jonathan D. Wren, Leonid Peshkin, Marko E. Horb

## Abstract

The diploid anuran, *Xenopus tropicalis*, has emerged as a key research model in cell and developmental biology. To enhance the usefulness of this species, we developed methods for generating immortal cell lines from Nigerian strain (NXR_1018, RRID:SCR_013731) *X. tropicalis* embryos. We generated 14 cell lines that were propagated for several months. We selected four morphologically distinct lines, XTN-6, XTN-8, XTN-10, and XTN-12 for further characterization. Karyotype analysis revealed that three of the lines, XTN-8, XTN-10, and XTN-12 were primarily diploid. XTN-6 cultures showed a consistent mixed population of diploid cells, cells with chromosome 8 trisomy, and cells containing a tetraploid content of chromosomes. The lines were propagated using conventional culture methods as adherent cultures at 30°C in a simple, diluted L-15 medium containing fetal bovine serum without use of a high CO_2_ incubator. Transcriptome analysis indicated that the four lines were distinct lineages. These methods will be useful in the generation of cell lines from normal and mutant strains of *X. tropicalis* as well as other species of *Xenopus*.

## Introduction

The speed and economy of modern genome, epigenome, transcriptome, and proteomic analyses provide unparalleled opportunity to benefit from specific experimental advantages of existing and emerging model organisms. For Xenopus laevis and Xenopus tropicalis, perhaps the most common amphibian model species, advantages include large clutches of embryos and external development making these organisms powerful systems for study of vertebrate development and physiology. Xenopus share with other vertebrates a large percentage of their genes and large regions of synteny with humans [1]. Inbred lines of animals of the two species have been developed and their genomes sequenced and annotated (www.xenbase.org). Of the two species in most common use, *Xenopus laevis* is more widely known. The use of *X. laevis* for genetic manipulations is somewhat complicated by the fact that it is an allotetraploid species, evolutionarily derived from the crossing of two related, ancestral species. As a result, many genes (approximately 56%) are expressed from two sets of alleles [2]. In contrast, *Xenopus tropicalis*, is a true diploid organism. Detailed developmental analyses of gene expression during early *Xenopus* development are available [3, 4]. Modern approaches using TALEN and CRISPR technologies have been used successfully to edit genes in embryos of both *X. laevis* and *X. tropicalis* [5-8].

Cell lines have several advantages for exploring biochemical pathways, including simplifying molecular analyses, simple physiological manipulation through application of drugs, and accessibility for high resolution microscopic imaging. Specifically, the tractability of Xenopus cell culture makes Xenopus cell lines ideal for research and teaching laboratories with minimal equipment and expense. Typically, Xenopus cell cultures are grown in simple incubators or at room temperature using a culture medium (70% L15 medium, 20% H2O and 10% serum). L15 medium does not require an enriched CO2 atmosphere. *Xenopus* cells will grow robustly in a desk drawer. Like any tissue culture, propagation of Xenopus cell lines requires sterile technique, but since they are unlikely to harbor pathogens of danger to humans, they do not require BSL2 precautions.

Over twenty cell lines derived from *X. laevis* adult or tadpole tissues are listed in the Cellosaurus database (https://web.expasy.org/cellosaurus). X-C, XTC, A6, XL177, XL2, XL58, and XLK-WG. X-C cells and XL177 were reported to be aneuploid [9, 10]. XL2 cells were described to contain two cell populations with chromosome numbers of 36 (the euploid number in X. laevis) and 74 [11]. Chromosome numbers in other X. laevis cell lines have not been reported. We have studied mechanisms of cell division and mitotic cell cycle control in cells in culture using the X. laevis S3 cell line [12]. This line was developed in the laboratory of Dr. Douglas Desimone at the University of Virginia. Chromosome spreads prepared from S3 cells show normal ploidy (unpublished observations). For X. tropicalis (euploid chromosome content of 20), only one permanent cell line has been described. This line, termed Speedy, has a stable aneuploid karyotype of 21 chromosomes being trisomic for chromosome 10 [13]. Here we describe the development and characterization of four novel cell lines, three with normal ploidy, developed from embryos from the inbred Nigerian strain of *X. tropicalis*.

## Methods

### Establishment of Primary Cultures

*Xenopus tropicalis* Nigerian strain (NXR_1018, RRID:SCR_013731) embryos were produced by in vitro fertilization then dejellied with cysteine using standard protocols [14]. Embryos were reared in shallow dishes in 0.1 X Marc’s Modified Ringer’s Solution (MMR) at 30 degrees. Twenty healthy stage 19 embryos were transferred to a 1.5 ml sterile microfuge tube, moved to a horizontal laminar flow hood, and rinsed 5 times with 1.5 ml of 0.1 X MMR containing gentamycin (final concentration 5µg/ml from 10mg/ml stock, Gibco), penicillin/streptomycin (1X from 100X stock, Mediatech), Fungizone (final concentration 0.5µg/ml from 250µg/ml stock, Gibco). Embryos were then rinsed once with sterile Phosphate Buffered Saline containing the same concentrations of gentamycin, penicillin/streptomycin and fungizone (PBS+). Embryos, still in their vitelline envelopes, were then treated briefly, approximately 10 – 15 seconds, with 70% ETOH with 3 to 4 inversions of the tube. The 70% ETOH was quickly removed and replaced with PBS+. Then the embryos were rinsed four times with PBS+.

After the last rinse, the PBS+ was replaced with Trypsin-EDTA (final concentration 0.25 X from 10 X stock, Atlanta Biologicals). The embryos were disrupted by gentle pipetting, and flicking of the tube. Too vigorous pipetting at this stage resulted in unacceptable levels of cell disruption. The embryos were incubated in the Trypsin-EDTA solution for 4 min with occasional inversion of the tube. Tubes containing the cells were centrifuged at 240 X g for two min to pellet most cells but leave yolk platelets released from broken cells in the supernatant. The supernatant was removed, and the cells were gently resuspended in Xenopus cell culture medium (70% calcium-free L15 medium (U.S. Biologicals, cat L2101-02 Leibovitz L-15 Medium w/L-Glutamine, Calcium-Free (Powder)), 20% H_2_O, 10% fetal bovine serum with 1 X penicillin/streptomycin). Tubes were centrifuged once more at 240 X g for 2 min, the supernatant discarded, and the cells resuspended in culture medium, then plated into a 60 mm tissue culture dish with low calcium L15-3G medium.

L15-3G medium was prepared as follows. Conditioned medium was prepared by growing cells from the *Xenopus laevis* S3 cell line in 75 cm^2^ tissue culture flasks. After 2 – 3 days of culture, medium was collected from these cells and centrifuged at room temperature on a clinical centrifuge at 520 X g to pellet debris. Originally, the medium was then collected into a sterile bottle and kept at 4°C. Subsequently, adding the medium to a bottle stored at −20°C was found to preserve growth promoting activity for longer times. When 200 – 400 ml of conditioned medium had been collected, it was thawed and sterile filtered through a 0.2 µm filter using vacuum filtration. The sterile medium was distributed to 50 ml sterile tubes (40 ml per tube) and placed at −20°C till use. When needed a tube was thawed, centrifuged at 520 X g to pellet any precipitates formed during freezing and thawing. The conditioned medium was mixed 1:1 with fresh medium and supplemented with 2 ng/ml basic fibroblast growth factor and 20 ng/ml epidermal growth factor. This medium was termed L15-3G and stored at 4°C.

After initial plating of the cells, dishes were sealed with plate sealing tape then incubated overnight at 30°C to allow viable cells to attach and spread. The next day the medium was removed, the dish rinsed gently with PBS to remove unattached cells and yolk platelets released from lysed cells. Fresh L15-3G medium was added to the dish, and the dish was resealed with tape and returned to the incubator. The culture medium was exchanged two times per week.

### Growth and Cloning of Cell Lines

Cells were allowed to proliferate until the dish was 80% - 90% confluent. For passage, cells were resuspended with full strength Trypsin-EDTA. To generate clones, cells were plated at very low density (∼100 cells) in 100mm tissue culture dishes. Cultures were observed daily to identify clones of proliferating cells derived from single progenitor cells. Clones were marked on the bottom of the dish with a diamond scribe objective and then circled on the bottom of the dish with a marker. Once colonies had grown to size of approximately 500 – 1000 cells, they were individually treated with Trypsin-EDTA and transferred to a 48 well tissue culture dish. To accomplish this, medium was withdrawn from the 100 mm plate and the plate was rinsed once with PBS. The PBS was removed and sterile, stainless-steel washers with 4 mm openings were placed over each colony. Trypsin-EDTA was added to the center of each washer and plates were observed until cells had rounded. At that point most of the Trypsin-EDTA was carefully removed, and the cells were suspended in 30 µl of L15-3G medium. The cells were then transferred to wells of a 48 well dish and fed with L15-3G medium. The outermost wells of the 48 well dish were not used for cells but were filled with sterile water as were any other extra wells not used for cells. Water-filled wells helped to minimize evaporation of medium in wells containing cells. The plates were sealed with plate sealing tape then placed at 30°C. Wells with successfully growing cultures were transferred to 24 well dishes, then 6 well dishes, to 25 cm^2^ flasks and then to 75 cm^2^ flasks. The flasks used were plug sealed and kept securely closed to prevent evaporation. When cell lines were growing vigorously, the L15-3G medium was replaced with Xenopus low calcium L15 medium with 10% FBS and pen/strep that did not include conditioned medium. A protocol for cell line maintenance is provided as a Supplemental file (see *Xenopus tropicalis* Cell Line Maintenance Protocols).

### Karyotype Analysis

Cell cultures at 60 – 70% confluency in 75 cm^2^ flasks were treated with 0.5 uM PD-166285 (inhibitor of Myt1 and Wee1 kinases) and 100 ng/ml Nocodazole (microtubule depolymerizer) for 3 h to enrich the mitotic cell population. Cultures were then treated with Trypsin-EDTA to release the cells from the culture flask, resuspended in 5 ml culture medium and centrifuged at 200 x g for 4 min. Cells were resuspended in 1 ml of culture medium. 500 ul of the cell suspension was distributed to two 1.5 ml microfuge tubes and centrifuged at 200 x g for 3 min. The medium was aspirated and cells were resuspended in swelling buffer, which consisted of 60% deionized H_2_O, 40% medium. Cells were incubated at 25 degrees Celsius for 15 – 20 min. Tubes were gently inverted several times to resuspend any cells that had settled. One ml of freshly made 3:1 methanol:acetic acid fixative was added dropwise to each tube, which was then kept undisturbed at room temperature for 15 min. Cells were centrifuged at 200 x g for 5 min. The fixative was aspirated, and the cells resuspended in 1 ml fixative. Cells were again centrifuged at 200 x g for 5 min. All but 30 – 40 ul of fixative was removed, and the cells were resuspended in the residual fixative by flicking the bottom of the tube. One hundred ul of fixative was then added to each tube and the cells were mixed using a 200 ul pipettor fitted with a wide bore tip.

Ethanol-washed 22 mm coverslips were placed at the bottom of a 15 cm petri dish on top of wetted filter paper. One to two drops of cell suspension in fixative were dropped onto each coverslip from a height of approximately 45 cm. After each coverslip was used, it was immediately transferred to the bench top on top of H_2_O-wetted kimwipes, and the coverslips allowed to dry for one hour or more. Coverslips were transferred to 6 well culture dishes and rehydrated with MilliQ water and labeled in H_2_O for 5 min with Dapi (Sigma-Aldrich, Cat # 9542) 1:10,000 dilution of 1 mg/ml aqueous stock and Syber Gold 1:20,000 dilution of DMSO stock (ThermoFisher Cat # S11494). The coverslips were then rinsed with H_2_0 3 times for 4 min each. The coverslips were then mounted on slides with 8 ul of Vectashield and the edges sealed with clear nail polish. Slides were observed and imaged with an Axioplan II microscope using a 100x 1.4 NA objective, an ORCA ER camera and Metamorph imaging software.

### Growth Analysis and Plating Efficiency

Cells were seeded in multiple wells of 6 well dishes and grown in growth medium at 30°C. Cells were seeded on day 0 and grown till day 7. Each day, with the exception of day 5, one well was trypsinized, and the cells were counted by hemacytometer. To test plating efficiency, cells were trypsinized, seeded at initial densities of 500, 1000 and 1500 cells per well in 6 well dishes, and incubated for 10 days. On day ten the cells were rinsed briefly in PBS then stained with 2% Methylene blue in 50% ethanol. They were washed 2x with water and visible colonies were counted.

For analysis of expression of senescence-associated beta galactosidase, XTN-12 cells were plated on 25 mm coverslips in 6 well dishes at approximately 1000 cells per well. The cells were cultured for 7 days then processed for beta-galactosidase expression using a Senescence Detection Kit (BioVision cat # K320-250) according to the manufacturer’s directions.

### RNA Sequencing and Analysis

For RNA isolation cells were grown in T75 flasks until 70% - 80% confluent, then trypsinized to create a single cell suspension. RNA was isolated using an RNeasy Mini Kit (Qiagen, # 74134) according to the manufacturer’s directions. Prior to RNA-seq analysis quality control measures were implemented as described previously [15]. The concentration of RNA was ascertained via fluorometric analysis on a Thermo Fisher Qubit fluorometer. The overall quality of RNA was verified using an Agilent Tapestation instrument. Following initial quality control steps, sequencing libraries were generated using the Illumina Truseq Stranded mRNA with Library prep kit according to the manufacturer’s protocol. Briefly, mature mRNA was enriched via pull down with beads coated with oligo-dT homopolymers. The mRNA molecules were then chemically fragmented, and the first strand of cDNA was generated using random primers. Following Rnase digestion, the second strand of cDNA was generated replacing dTTP in the reaction mix with dUTP. Double stranded cDNA then underwent adenylation of 3’ ends following ligation of Illumina-specific adapter sequences. Subsequent PCR enrichment of ligated products further selected for those strands not incorporating dUTP, leading to strand-specific sequencing libraries. Final libraries for each sample were assayed on the Agilent Tapestation for appropriate size and quantity. These libraries were then pooled in equimolar amounts as ascertained via fluorometric analyses. Final pools were absolutely quantified using qPCR on a Roche LightCycler 480 instrument with Kapa Biosystems Illumina Library Quantification reagents.

For RNA-seq, paired-end 75bp read sequencing was performed at 4 time points on an Illumina NextSeq 500 sequencing platform. Raw reads, in a FASTQ format, were trimmed of residual adaptor sequences using the Scythe software. Low quality bases at the beginning and end of reads were removed using Sickle, then the quality of remaining sequences was confirmed with FastQC. Further processing of quality sequencing reads was performed with utilities provided by the Tuxedo Suite software. Reads were aligned to the Xenopus tropicalis v9.0 genome reference using the TopHat component.

## Results

### Pitfalls in establishing stable cell lines from Xenopus tropicalis embryos

One of the most significant difficulties in establishing stable cell lines from Xenopus embryos is creating and maintaining stringent sterile conditions for the cultured cells. Surface sterilization procedures can be applied during the dejellying process [14]. However, these are generally insufficient to guarantee sterility within cultures of the dissociated embryo cells. Therefore, our protocols emphasize the use of antibiotics during dissociation and, just before dissociation, treatment of embryos within the intact fertilization envelope with 70% ethanol, all within the confines of a sterile culture hood. These precautions are usually, though not always, sufficient to guarantee sterility. Fungal contaminants do occasionally occur but are not common enough to necessitate routine use of fungicides.

A second difficulty encountered during embryo dissociation is disruption of cells resulting in release of numerous yolk platelets that can restrict cell attachment to the culture surface. As noted in the methods, at the stage of embryo dissociation, vigorous pipetting is to be avoided. Contamination of the cultures by released yolk platelets is minimized by differential centrifugation steps. If large numbers of free yolk platelets remain after cells have attached to the substratum, they can be removed the next day by gentle pipetting, discarding the released platelets with the medium and refreshing the culture with fresh medium. Initially many attached cells will contain internal yolk platelets. These will decrease over time in culture and eventually disappear.

Once cells attach, they will begin to proliferate. However, if cells are plated at too low a density, many will undergo a terminal differentiation or senescence. This can be reduced but not eliminated by the use of conditioned medium. Thus, in the initial stages it is beneficial to maintain primary cultures at high density, preferably with 25% to 50% of the substrate occupied by cells. Then when the primary cultures become near confluent, transfer no less than 25% of the resuspended cells to new culture flasks. Eventually robust cell growth will occur and cells can be passaged at greater dilution and even cloned though not all cells plated at very low concentration for cloning purposes will generate successful colonies.

### Xenopus tropicalis cell lines

We established 14 cloned cell lines that were passaged in culture for over 6 months. Some of the lines appeared morphologically similar suggesting that they were multiple isolates of identical or closely related cells. Based on distinct morphologies, we selected 4 lines for more detailed analysis (Figure 1). We named these cell lines with the prefix XTN, which stands for *<u>X</u>enopus tropicalis <u>N</u>igerian* strain. The lines selected were named XTN-6, XTN-8, XTN-10 and XTN-12.

**Figure 1.**
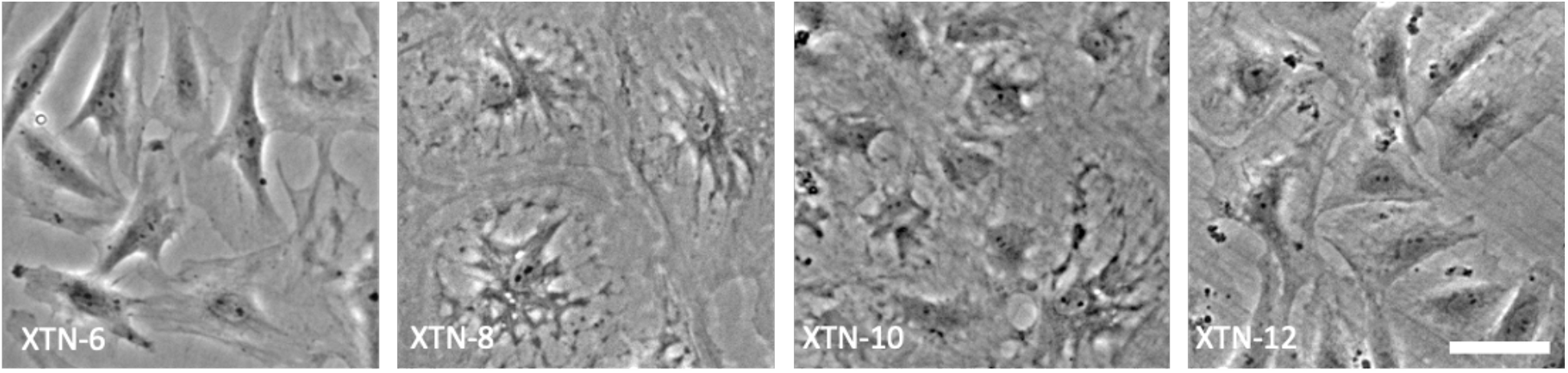
Phase contrast images of four selected Xenopus tropicalis cell lines. Bar = 50µm.

### Karyotype analysis

Chromosome spreads prepared from the four selected *X. tropicalis* cell lines revealed predominantly diploid karyotypes of 20 chromosomes for XTN-8, XTN-10 and XTN-12 cell lines while cells from the XTN-6 line revealed more complex features (Figure 2). Most XTN-6 cells contained 21 chromosomes although a significant number showed the normal complement of 20. Chromosome spreads of XTN-6 cells also revealed a large percentage (41.3%) of the cells with more than 21 chromosomes were tetraploid containing 40 to 42 chromosomes. Detailed analysis revealed that the extra chromosome in XTN-6 cells with 21 chromosomes was not a random aneuploidy but reflected a conserved trisomy of chromosome 8.

**Figure 2.**
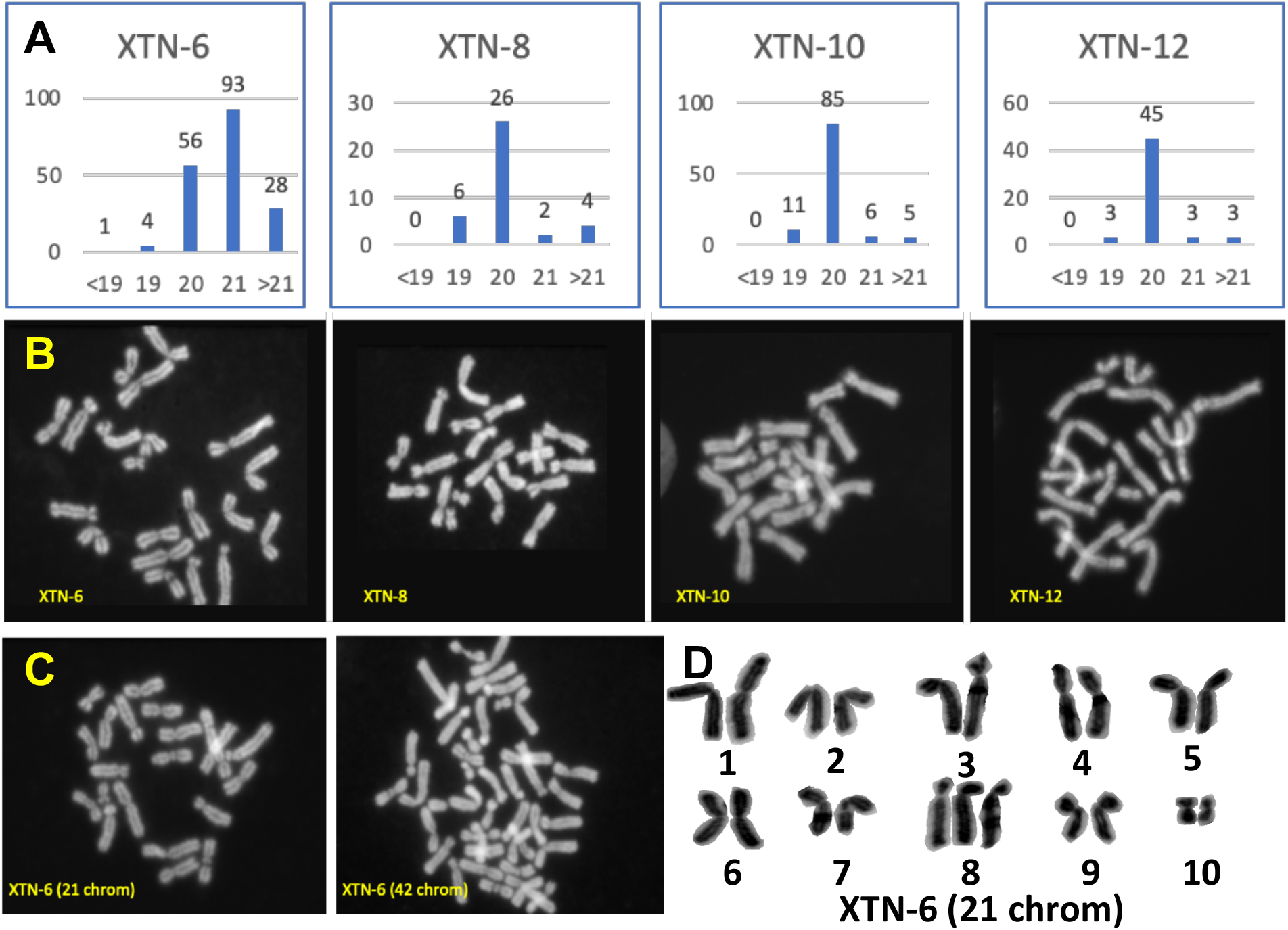
Karyotype analysis of four selected X. tropicalis cell lines. A) Quantitation of chromosome spreads from XTN-6, XTN-8, XTN-10, and XTN-12 cell lines. Karyotype categories are listed on the X axis and numbers of spreads of each category are listed above the bars. B) Examples of normal diploid spreads (20 chromosomes) from each cell line. C) Chromosome spreads from XTN-6 cells showing single examples of 21 chromosomes, single chromosome aneuploidy and 42 chromosomes, possibly a tetraploid cell derived from a cell which originally contained 21 chromosomes. D) An example of a karyotype analysis which revealed that XTN-6 cells with 21 chromosomes contain an extra copy of chromosome 8. (Chromosomes are numbered according to the updated nomenclature [16]).

### Growth characteristics

Cells seeded in a six well dish at day 0 grew exponentially until day 6 when they began to plateau (Figure 3). During exponential growth XTN-6, XTN-8, and XTN-10 cells showed a doubling time of approximately 24 hours at 30°C. XTN-12 cells grew slightly faster with a doubling time of about 22 hours. We also tested the plating efficiency, namely their ability to form colonies when cells were plated at low density in six well dishes. Generally, cloning efficiency was highest in XTN-8 and XTN-10 cell lines followed by XTN-12 with XTN-6 cells being the least efficient (Table 1)

**Table 1.** Plating efficiency of four selected X. tropicalis cell lines seeded at low density in 6 well dishes.

| Number of Cells Plated | 500 | 1000 | 1500 |
| --- | --- | --- | --- |
| XTN-6 colonies formed | 6 | 14 | 23 |
| XTN-8 colonies formed | 98 | 169 | 207 |
| XTN-10 colonies formed | 24 | 108 | 157 |
| XTN-12 colonies formed | 28 | 70 | 96 |

**Figure 3.**
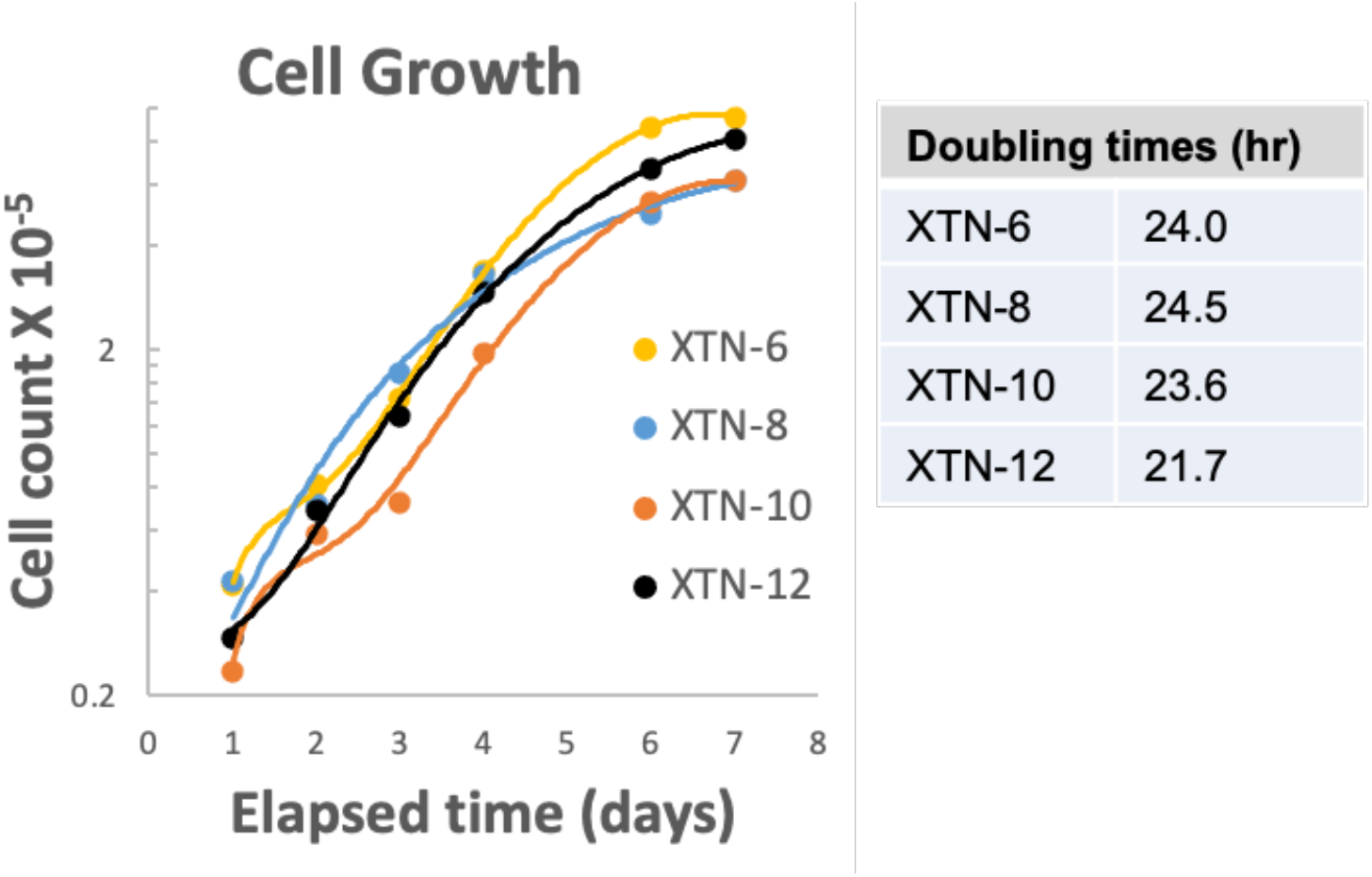
Growth properties of four X. tropicalis cell lines.

To examine whether plating efficiency might be hampered by a tendency of cells at low density to undergo senescence, we tested one of the lower efficiency lines, XTN-12, for expression of beta-galactosidase, one of the most common senescence markers [17]. XTN-12 cells plated at low density on coverslips and cultured for 7 days showed cells that were positive for senescence-associated beta-galactosidase, consistent with the interpretation that they had undergone senescence-dependent cell cycle arrest (Figure 4).

**Figure 4.**
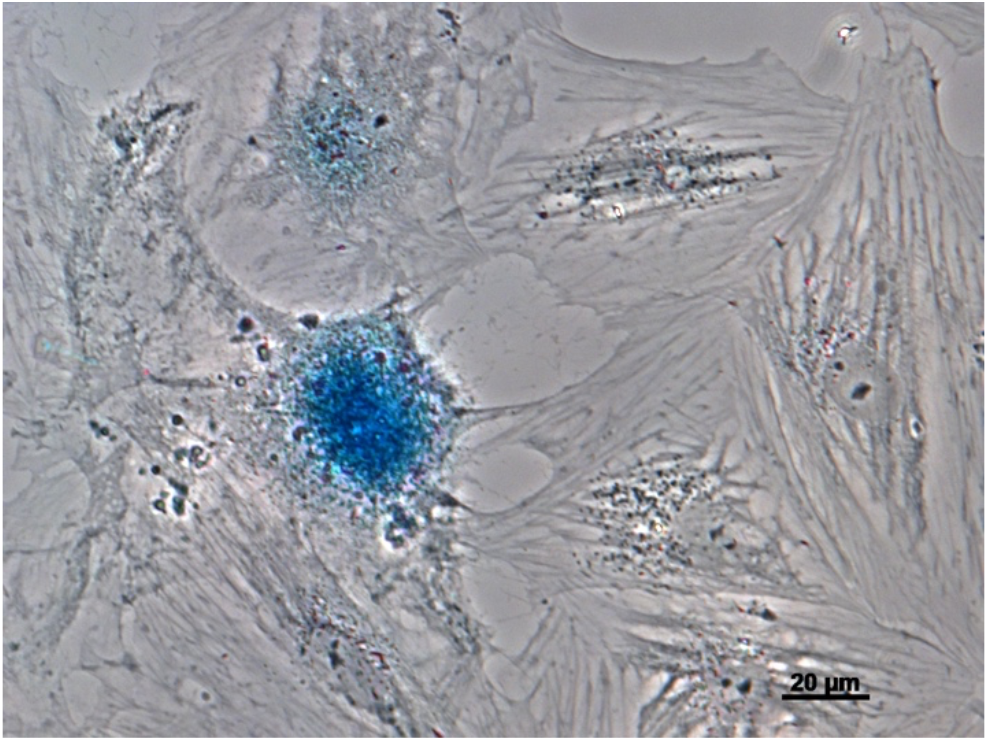
Expression of the senesce marker beta-galactosidase indicated by the blue color in this phase contrast image of XTN-12 cells initially plated at low density.

### Transcriptome Analysis

Consistent with morphological differences observed among the four cell lines, gene expression profiles revealed considerable dissimilarities among the lines (Figure 5A and Supplemental Table 1). As indicated in the dendrogram (Figure 5B) lines XTN-8 and XTN-10 were most similar to each other and XTN-6 the most distinct.

**Figure 5.**
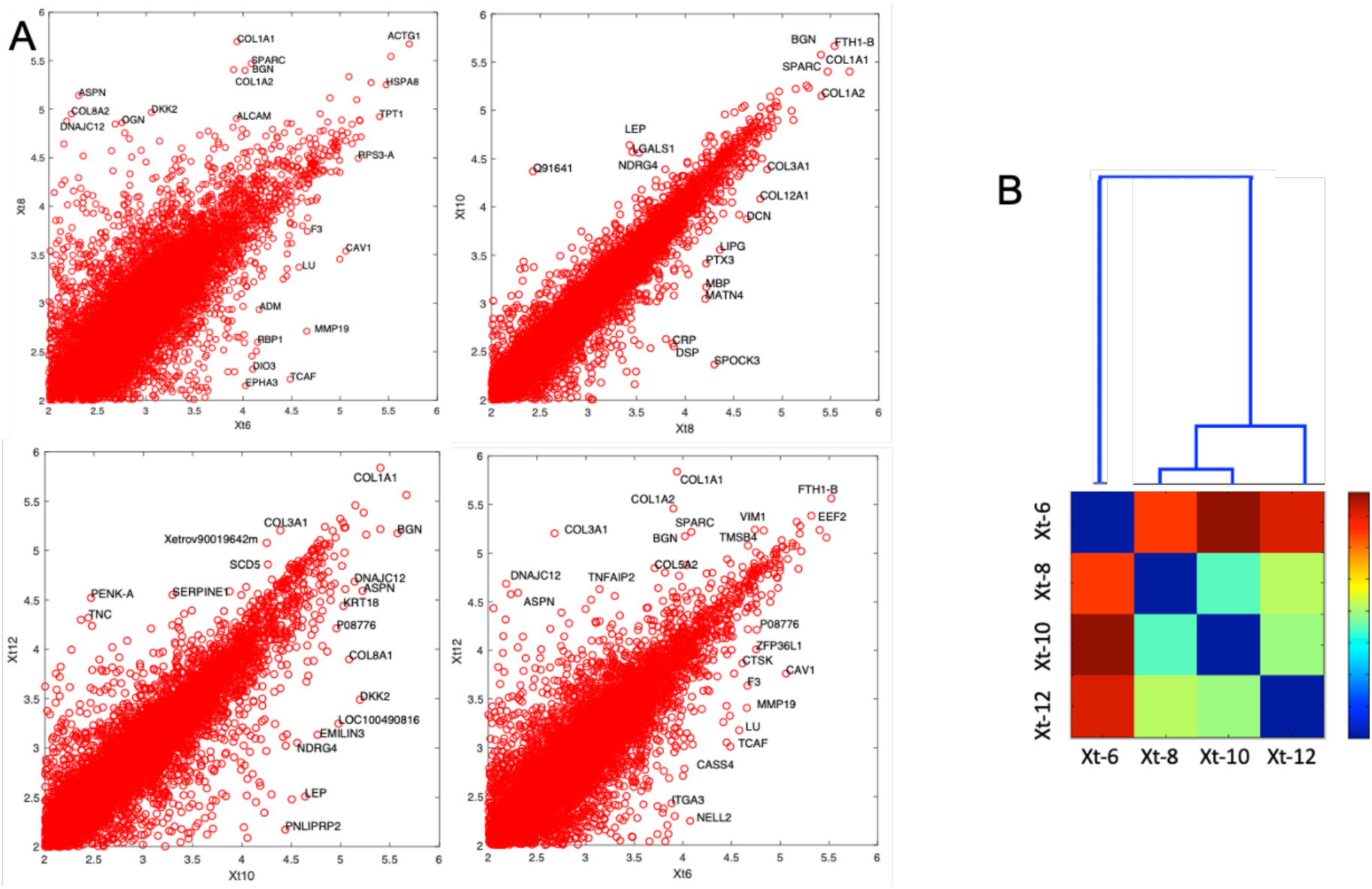
A) Four pair-wise comparisons of gene expression in cell lines. The most highly expressed genes and outliers are labeled using a human symbol of a homologous gene. It is apparent that XTN-8 and XTN-10 are most similar to each other and XTN-6 is most different from the rest. B) Dendrogram and expression comparing all four cell lines. Red indicates lowest similarity and blue highest.

In order to characterize our cell lines further, we compared transcriptomics gene expression profiles to the single cell expression data from the Xenopus tropicalis embryos profiled around the same developmental stage as we derived the lines from [3]. The bottom-up bulk analysis of entire transcriptome does not single out any of approximately 200 groups clustered by cell type in that paper (data not shown). We then took a top-down approach focusing on the key marker genes curated from the single cell data [3] and from Xenopus Anatomy Ontology available at Xenbase https://www.xenbase.org/anatomy/xao.do?method=display&tabId=0). This is presented in Supplemental Table S2 which lists 50 marker genes showing strong differences in expression among the cell lines. While we again do not see a clear indication of the cell lines expressing a pattern characteristic to just one of the embryonic cell lineages, there are some features that stand out. First, XTN-6 is again the most distinct from the other lines. XTN-6 shows strong differential expression of individual gene markers characteristic of several cell types, as different endoderm (*gjb2*) and intermediate mesoderm (*osr2*), while the other strong markers for these same cell types are detected in different cell lines - e.g. in XTN-10 for endoderm marker (*c8g*) and XTN-8 for intermediate mesoderm (*tbx1*). Yet, XTN-6 shows consistent expression of somite progenitor genes exclusively expressing many markers: *hoxc10, hoxc11, hoxa11, hoxd1, hoxd4, sp5*.

XTN-8, XTN-10, and XTN-12 show expression of several markers typical for pre-placodal ectoderm (*pitx2, nkx2-3, olfm4, six1, eya2*). XTN-12 over-expresses collagen 9 isoforms (*col9a1, col9a2 and col9a3*) typical for notochord. All four cell lines show characteristics of the endoderm and the gene program typical for migrating cells (*fli1, snai2, twist1, sox8, sox9*). XTN-6 shows enhanced expression of genes located on chromosome 8, as might be expected given the high proportion of cells trisomic for this chromosome. (Note that this chromosome is annotated as chromosome 10 in Xenbase, consistent with the older chromosome numbering system used for *X. tropicalis* [18].)

## Discussion

Elucidating molecular pathways, mapping gene regulators and studying the consequences of mutation and drug treatments is often much simpler through use of cell lines in mammalian research. Such advantages hold true for *Xenopus* research as well but cell line resources have been limited, particularly for the diploid *X. tropicalis*. Here we have developed four new cell lines, three of which are normal ploidy. Morphology and RNAseq differences reveal that the cell lines are likely derived from distinct founder cells from the embryo. Our methods for developing new *Xenopus* cell lines are relatively simple and could be applied to develop lines from the embryos of currently available mutant *Xenopus* animals.

The sexual maturation time of X. laevis and X. tropicalis are ∼6-12 months, respectively, making their use in standard genetic crosses problematic. In addition, there is no method for long term storage of frog eggs, embryos, or adults. Frogs must be kept as live animals to maintain a genetic line. Although success has been achieved in freezing Xenopus sperm, successful regeneration of a mutant animal line, even under the best circumstances, requires 1-2 years since two generations must be bred. Finally, while transgenesis and gene deletion by TALENs or CRISPR technology is established in Xenopus, their current application to fertilized Xenopus eggs [3-5] and to Xenopus oocytes [2, 6, 7] generates mosaic embryos and does not allow for selection of specific, defined mutations or gene insertions.

By using Xenopus cell lines as the basic unit of genetic manipulation, as opposed to the current methods of embryo injection, specific gene manipulations can be selected in cell clones and fully characterized at the molecular level, prior to generation of the mutant animal. Mutant cell lines are easily frozen and can be stored indefinitely, at low cost, as frozen stocks. Potentially, somatic cell nuclear transfer will allow the production of mutant frogs in the first (F0) generation, negating the need for breeding. Even mutations that might lead to infertility or embryonic lethality can be studied in the F0 generation. However, nuclear transfer efficiency from current cell lines is inefficient (data not shown). This is likely due to difficulty in epigenetic reprogramming of the transferred nuclei. Potentially, the development of induced pluripotent stem cells from the cell lines will greatly increase the efficiency of embryo production after nuclear transplantation.

## Supporting information

Supplemental Table 1 - RNAseq expression data for cell lines

Supplemental Table 2 - Differentially expressed marker genes

## Acknowledgements

This work was supported by Whitman fellowships to GJG from the Marine Biological Laboratory, by grant # 1645105 to GJG and MEH from the National Science Foundation. LP has been supported by grant # R01HD073104 from the National Institute of Child Health and Development.

## Supplemental Material

### Xenopus tropicalis Cell Line Maintenance Protocols

#### Solutions

- 10X Trypsin EDTA Atlanta biological cat# B81210, probably source does not matter
  - 0.85% PBS (use to dilute trypsin-EDTA to 1X) Store in fridge or at −20°C (normal strength PBS is also usable)
- 70% complete calcium-free L-15 (70% L15, 10% FBS, 20% H_2_O)
  - powder calcium free L-15 - US biologicals #L2101-02 powder, smallest amount that can be purchased is powder for 50 Liters
  - For 70% (amphibian medium) measure 9.63 g into 800 ml water in flask with stir bar and stir for 10 min till fully dissolved
  - add water to total 900 ml (other 100 ml will be serum)
  - sterilize by filtration and separate into two sterile bottles with 450 ml each
  - keep at 4 degrees, just before using add 50 ml sterile FBS
  - from 100x concentrate, add Pen/Strep to normal mammalian cell final concentration

#### Maintaining Xenopus cell cultures

- Cells are grown in standard tissue culture flasks or dishes. To avoid evaporation, use plug seal flasks and close tightly or use plate sealing tape on culture dishes. (We do not use humidified incubators for our *Xenopus* cell cultures.)
- Typically we grow *X. tropicalis* cells at 30°C. However, cells will also grow, albeit more slowly at room temperature.
- L15 medium should be used with normal atmosphere, NOT with high CO_2_.
- To passage cells
  - Remove medium by aspiration.
  - Add 1x trypsin-EDTA, immediately aspirate to remove.
  - Add second aliquot of 1x trypsin-EDTA, incubate until cells start to round up. Time varies among different lines.
  - When cells are mostly rounded carefully remove almost all trypsin without dislodging cells.
  - Wait until cells are fully rounded. Test release by banging flask on edge to see if cells release from the flask.
  - Add 2 – 5 ml medium.
  - Resuspend by pipetting with sterile transfer pipette or sterile graduated pipette.
  - Transfer to new flask or dish. probably dilute no more than 1:4 initially and determine best dilution empirically for each line.*

#### Storing cell cultures in liquid nitrogen

- When cells are nearly confluent, resuspend cells as above in complete medium. Have a final volume of about 2 ml from a small (T25) flask or about 5ml from a large (T75) flask.
- Put cells in 15 ml polypropylene tube.
- Rapidly pipette in DMSO to 10% final concentration and flick tube to mix.
- Aliquot 1 ml to cryotubes.
- Place between 2 styrofoam racks from 15 ml tubes or commercial cryocooler.
- Place at −80 overnight.
- Transfer vials to liquid nitrogen tank.

#### Thawing frozen cells

- Swish vial in 37°C bath until it just starts to thaw (don’t let cells get to 37°C)
- Transfer cells to T25 flask (or T12.5 flask if they are expected to be dilute).*
- Add 3 – 5 ml of medium.
- Allow cells to attach overnight.
- The next day change medium to remove DMSO and dead cells.

*In some lines, cells will tend to senesce if cultured at too low a concentration. However, lines can be cloned in the presence of conditioned medium.

